# Formation of polyphasic RNP granules by intrinsically disordered Qβ coat proteins and hairpin-containing RNA

**DOI:** 10.1101/2024.10.26.620452

**Authors:** Naor Granik, Sarah Goldberg, Roee Amit

## Abstract

RNA-protein (RNP) granules are fundamental components in mammalian cells where they perform multiple crucial functions. Many RNP granules form via phase separation driven by protein-protein, protein-RNA, and RNA-RNA interactions. Notably, associated proteins frequently contain intrinsically disordered regions (IDRs) which can associate with multiple partners. Previously we have shown that synthetic RNA molecules containing multiple hairpin coat-protein binding sites can phase separate, forming granules capable of selectively incorporating proteins inside. Here, we expand this platform by introducing a phage coat protein with a known IDR which facilitates protein-protein interactions. We show that the coat protein phase-separates on its own *in vivo*, and that introduction of hairpin-containing RNA molecules can lead to dissolvement of the protein granules. We further demonstrate via multiple assays that RNA valency, determined by the number of hairpins present on the RNA, leads to distinctly different phase behaviors, effectively forming a polyphasic programmable RNP granule. Moreover, by incorporating the gene for a blue fluorescent protein into the RNA, we demonstrate a phase-dependent boost to protein titer. These insights not only shed light on the behavior of natural granules, but also hold profound implications for the biotechnology field, offering a blueprint for engineering cellular compartments with tailored functionalities.

## Introduction

Biomolecular condensates are micron-scale, membrane-less compartments which are involved in numerous cellular processes^1,2^. These condensates, often referred to as membraneless organelles, are comprised of dozens to hundreds of macromolecules concentrated into small volumes. There is a growing recognition that many biomolecular condensates form through the phenomenon of phase separation, in which one or more components drive a demixing transition leading to the formation of at least two coexisting phases^3–6^. Of the many types of biomolecular condensates, ribonucleoprotein (RNP) granules have attracted significant interest from the scientific community^7–10^, primarily due to their involvement in gene expression^11^. Furthermore, research has linked disruption in RNP granule function to the onset of specific cancers and neurological conditions^12^. RNP granules are widespread in eukaryotic cells, with notable examples including the nucleolus, stress granules and P-bodies. Granules were also recently found to exist in bacterial cells, where they play a role in mRNA degradation^13^.

RNP granules are broadly characterized as macromolecular assemblies of RNAs and proteins, with distinct granules containing different sets of macromolecules. The phase separation process of these granules is facilitated through multivalent RNA-RNA, RNA-protein, or protein-protein interactions. Notably, RNA molecules play an active role in both the formation and dissolution of granules. This has been demonstrated in vitro, where RNA in small concentrations was shown to promote granule formation, while RNA in concentrations higher than a certain threshold led to granule dissolution^14,15^. In addition, it has been demonstrated that RNA can modify certain physical characteristics of granules, such as their three-dimensional structure, in a concentration-dependent manner^16–19^.

While RNAs play a crucial role in the formation and regulation of RNP granules, they represent only one aspect of the system as these granules cannot exist without a protein component. Key properties of granule-associated proteins include the existence of RNA-binding domains, and intrinsically disordered regions (IDRs). IDRs are protein segments which exhibit rapid conformational changes at the nano-to micro-second timescales, rendering them unlikely or incapable of forming a stable structure^20–23^. Due to their unstructured nature, IDRs possess remarkable flexibility in their ability to form protein-protein interactions varying over a wide range of valencies and a broad-range of interactions strengths^24–26^. This flexibility enables them to associate with many cellular partners for a variety of functions^27,28^. In the context of phase separation, IDRs have been consistently reported as being able to change the material properties of condensates, from solid-like to liquid-like, or vice versa^29–32^.

Given the complex mixture of proteins and various RNA molecules in RNP granules, it is necessary to establish a theoretical framework to study the contribution of each species. Polyphasic linkage theory, formulated by Wyman and Gill^33^, has been extensively developed and utilized to explain how binding and linkage relations control phase transitions^34,35^. This formalism considers multivalent molecules driving phase transitions as scaffolds, and molecules which bind the scaffolds as ligands. Ligands do not undergo phase separation on their own and are not required for the phase separation of the scaffolds. However, they bind preferentially to scaffolds across phase boundaries and can either promote or destabilize condensates in cells.

Recently, we have shown that synthetic RNP granules can be formed by mixing hairpin-forming RNA molecules together with phage coat proteins (CPs) which bind the RNA. Using super resolution microscopy, fluorescence microscopy, and spatiotemporal tracking, we concluded that the resulting particles are gel-like in nature and are formed via liquid-solid phase separation, both *in vivo* and *in vitro*.

In addition, by changing the number of hairpins encoded into the RNA sequence, we were able to alter the biophysical characteristics of the granules, effectively forming a programmable platform^36^.

As constructs designed to investigate RNA-based phase separation, our synthetic condensates lacked intrinsically disordered protein elements. We hypothesized that introducing an IDR to our system could lead to a different set of behaviors and provide another mechanism of control over synthetic granule characteristics and dynamics. Here, we increase the complexity of our synthetic RNP platform by introducing a phage coat protein with a well-documented disordered region (Qβ), allowing for multivalent interactions between the proteins. We demonstrate that the Qβ coat protein can phase separate on its own when overexpressed in cells, and that introduction of hairpin containing RNA effectively modulates this behavior, yielding a complex RNA-valency dependent phase behavior consistent with polyphasic linkage model predictions. We explore the behavior of the condensates as a function of RNA valency as determined by the number of hairpins encoded into the RNA and demonstrate that this platform can be utilized to drastically overexpress proteins in bacteria.

## Materials and Methods

### Construction of slncRNA plasmids

The sequences for Qβ-1x/3x/5x/10x/12x/17x/NS (available in the supplementary table) were generated using the CARBP online tool^37^ and ordered from Integrated DNA Technologies (IDT) as gBlock gene fragments downstream to a T7 promoter and flanked by ApaI and AvrII restriction sites. The gBlocks were cloned into a pPROLAR vector containing a kanamycin resistance gene, which was linearized using the above-mentioned restriction enzymes (New England Biolabs (NEB), catalog numbers: R0114S and R0174S, respectively), and excised from 1% agarose gel following electrophoresis to obtain the relevant DNA segment. Sequences were verified using Sanger sequencing.

The DNA sequence of the TagBFP protein (available in the supplementary table) was ordered from IDT as a gBlock gene fragment flanked by AvrII and AgeI restriction sites and ligated to the slncRNA-containing pPROLAR vectors linearized using the same restriction enzymes (NEB, catalog numbers: R0174S and R3552S, respectively).

### Design and construction of fusion-RBP plasmid

The sequence for the Qβ phage coat protein lacking a stop codon was ordered from GenScript, Inc., and cloned into an A133 plasmid backbone between restriction sites KpnI and AgeI, immediately upstream of an mCherry gene lacking a start codon, under the RhlR promoter containing the rhlAB las box and induced by N-butyryl-L-homoserine lactone (C4HSL) (Cayman Chemicals). The backbone contained an ampicillin (Amp) resistance gene.

### Fluorescence recovery after photobleaching (FRAP) measurements

BL21 (DE3) cells expressing the two-plasmid system (RNA expression plasmid, and QCP-mCherry or mCherry expression plasmid) were grown overnight in 5 ml LB, at 37°with appropriate antibiotics (Kan, Amp). Overnight culture was diluted 1:100 into 3 ml semi-poor medium consisting of 95% bioassay buffer (BA: for 1 L—0.5 g Tryptone [Bacto], 0.3 ml glycerol, 5.8 g NaCl, 50 ml 1 M MgSO_4_, 1 ml 10×PBS buffer pH 7.4, 950 ml DDW) and 5% LB supplemented with antibiotics, and induced with Isopropyl β-D-1-thiogalactopyranoside (IPTG) (final concentration 0.33 mM) and N-Butanoyl-L-homoserine lactone (C4HSL) (final concentration 50 μM). Culture was shaken for 4-6 hours at 37°C before measurements.

Gel slides for bacterial spatial fixation were prepared as follows: A solution of 1.5% (w/v) low melting agarose (Lonza) in PBS was heated for approximately 20 seconds to allow full melting of the agarose. Approximately 50 μl was deposited on a microscope slide, and a coverslip was immediately applied. When the agarose hardened, the coverslip was carefully removed, leaving a thin pad of agarose on the slide. Ten microliters of bacterial liquid culture were then deposited on a fresh coverslip, and the microscope slide and pad were placed on top with the agarose facing down so that the culture was spread between the pad and coverslip. Translucent nail polish was applied to the edges of the coverslip to prevent evaporation.

FRAP measurements were done using an LSM-880 laser scanning confocal microscope (Zeiss) with an sCMOS camera, and a 63x 1.46 NA oil immersion objective. Following identification of a live cell, photobleaching of a localized region was done using a 488 nm laser at full power (25 mW). mCherry fluorescence was measured using excitation with a 594 nm laser with 1% power (0.1 mW) with images being taken every 15 seconds for a total of 5 minutes. FRAP image analysis was done using the ImageJ software with the Stowers plugin suite.

### Structured illumination super resolution microscopy

Super resolution images were captured using the Elyra 7 eLS lattice SIM super resolution microscope (Zeiss) with an sCMOS camera, a 63×1.46 NA water immersion objective, with a 1.6x further optical magnification. 405 nm, and 585 nm lasers were used for excitation of the BFP, and mCherry, respectively. Microscope control and data acquisition was performed using ZEN Black version 3.3.89. 16-bit 2D image sets were collected with 13 phases and analysed using the SIM^2 image processing tool by Zeiss.

### Flow cytometry measurements

Flow cytometry measurements were performed using a MACSQuant VYB flow cytometer (Miltenyi Biotec), with the 561 nm excitation laser and the Y2 detector channel (a 615/20 nm filter). The flow cytometer was calibrated using MACSQuant calibration beads (Miltenyi Biotec) before measurement. Running buffer, washing solution, and storage solution were all purchased from the manufacturer (Miltenyi Biotec, catalog numbers 130092747, 130092749, and 130092748, respectively).

Events were defined using an FSC-height trigger of 3. An FSC-area over SSC-area gate (gate-1) was created around the densest population in the negative control (non-induced cells), and events falling inside this gate were considered live bacteria. From the gate-1-positive cells, an SSC-height over SSC-area gate (gate-2) was created along the main diagonal, and each event falling inside this gate was considered a single bacterium. Finally, from the gate-2-positive cells, a histogram of mCherry/BFP distribution was created, with the threshold for positive cells set by leaving around 0.1% positive events in the negative control.

For mCherry/QCP-mCherry measurements, IPTG and/or C4HSL-induced or non-induced *E. coli* BL21 cells were grown overnight at 37°C with appropriate antibiotics. In the morning, the bacterial culture was diluted 1:100 into 3 ml of semi-poor medium consisting of 95% BA and 5% LB, supplemented with the relevant inducers (IPTG and/or C4-HSL, or none). The diluted cells were shaken for six hours prior to measurement to allow sufficient time for induction and protein expression. Samples and appropriate controls were loaded onto a 96-well plate (ThermoFisher Scientific, catalog number: 167008) in triplicates, each well containing 200 μl of diluted bacterial cells.

For BFP measurements, IPTG and/or C4HSL-induced, or non-induced *E. coli* BL21 cells were grown overnight in 5 ml LB at 37°with appropriate antibiotics and relevant inducers. In the morning, the bacterial culture was centrifuged at 4000 RPM for 10 minutes and the liquid media was discarded. Cells were resuspended in 5 ml semi-poor medium consisting of 95% BA and 5% LB. Samples were loaded onto a 96-well plate in triplicates, each well containing 200 μl of diluted bacterial cells.

### Real time PCR measurements

Starters of E. coli BL21 containing the relevant constructs on plasmids were grown in 5 ml LB medium with appropriate antibiotics and inducers overnight. For each construct, three different colonies were selected as biological replicates. The following morning, cultures were pelleted by centrifugation at 5000g for 5 minutes and resuspended in 750 μl RNALater solution (ThermoFisher Scientific, catalog number AM7020). Samples were stored at 4°c until RNA extraction.

RNA was extracted using the NucleoSpin RNA kit (Macherey-Nagel, catalog number: 740955.50) according to the manufacturer’s protocol. Extracted RNA was treated with DNase to remove residual DNA using the Turbo DNase kit (ThermoFisher Scientific, catalog number: AM2238). Reverse transcription was done using the High-Capacity cDNA Reverse Transcription kit (Applied Biosystems, catalog number: 4368814).

Primer pairs for TagBFP and the housekeeping gene idnT were chosen using the Primer Express software. RT-PCR was performed on a QuantStudio 12K Flex machine (Applied Biosystems) using SYBR-Green reagents. Three technical replicates were measured for each of the three biological replicates. A Ct threshold of 0.2 was chosen for all genes.

## Results

### RNA with valency of n=3 inhibits formation of Qβ coat protein homotypic biomolecular condensates

Our previously described RNP granule platform utilized the PP7 bacteriophage coat protein (PCP) as the protein component^36^. PCP is characterized by a rigid tertiary structure containing multiple β-strands, rendering it highly ordered and thus lacking the unique characteristics of IDRs^38^. We opted to add an element of disorder while still retaining the simplistic two-component nature of our synthetic granules. Therefore, we decided to utilize the coat protein of the bacteriophage Qβ, which contains a documented disordered region^39^ and is identified as such by state-of-the-art disorder predictors^40^ (Figure 1A). We hypothesized that when overexpressed, the Qβ coat protein (QCP) would phase separate in cells on its own accord to form homotypic biomolecular condensates. To test for this, we fused QCP on its C-terminus to an mCherry fluorescent protein, both to tag the coat protein and to render it incapable of spontaneously forming the well-known viral capsid^41–44^. We cloned the fusion protein into *E. coli* cells under the control of an inducible pRhlr promoter and tested for the formation of bright polar puncta using a fluorescence microscope. Per our assumption, we found bright spots localized to the cell poles consistent with homotypic biomolecular condensates for most cells examined (Figure 1B), indicating that the disordered region within QCP provides sufficient multivalency to drive some form of phase separation.

**Figure 1.**
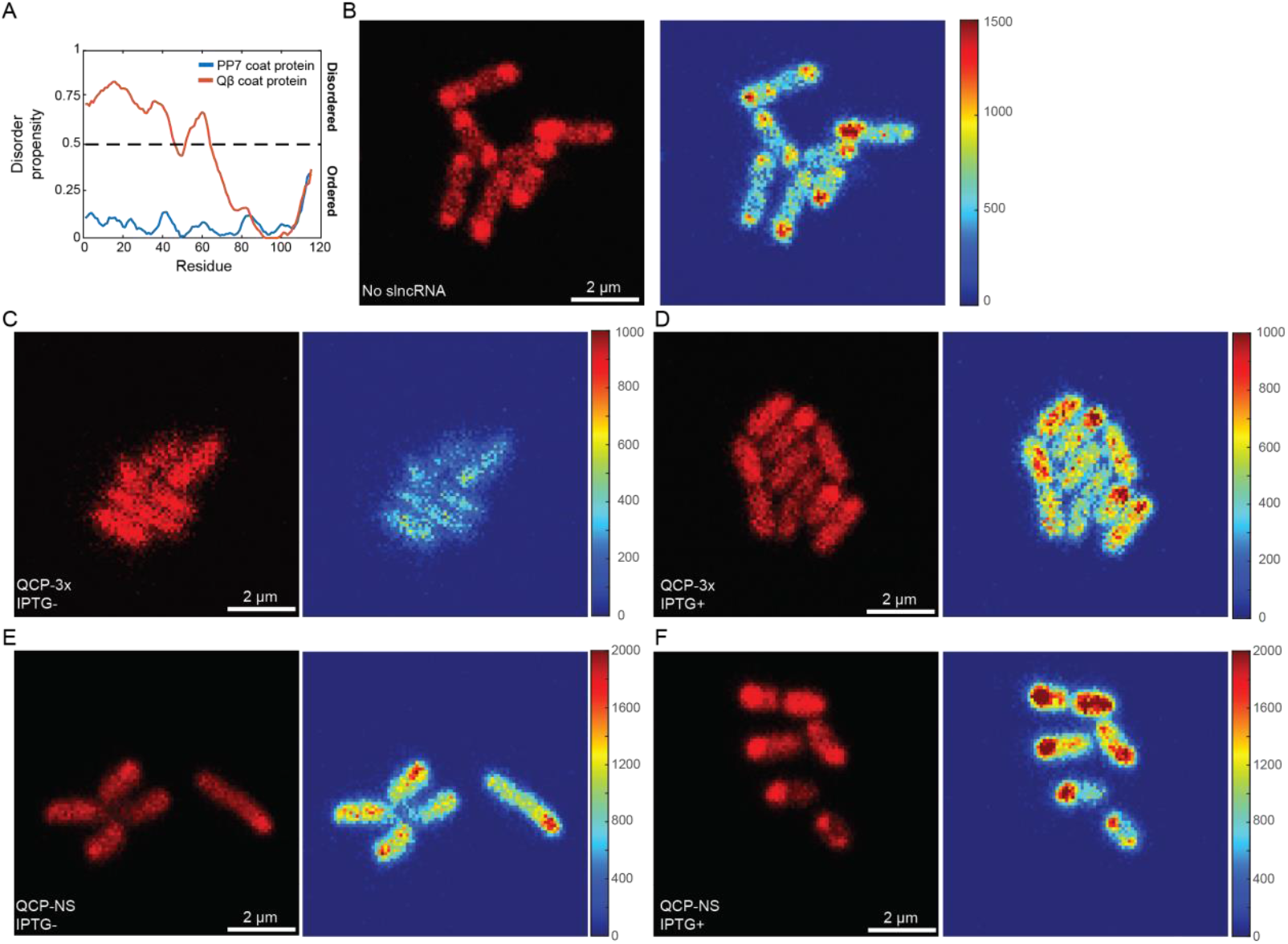
QCP phase separation behavior in the absence or presence of different slncRNAs. **A**. Disorder propensity predictions of the Qβ and PP7 phage coat proteins, in red and blue, respectively. The predictions show a high confidence in the existence of a disordered region in QCP. Prediction by Metapredict. **B**. Sample field of view showing cells expressing just the QCP-mCherry protein with no slncRNA. The cells exhibit bright fluorescent spots localized to the cell poles, reminiscent of phase separated compartments. **C-D**. Sample fields of view showing cells expressing the QCP-mCherry protein together with a slncRNA encoding for three hairpin binding sites, without slncRNA induction (**C**), and with it (**D**). Cells without induction exhibit uniform fluorescence across the cell with no discernible puncta. With slncRNA induction, bright fluorescent spots reappear at the cell poles. **E-F**. Sample fields of view showing cells expressing the QCP-mCherry protein together with NS slncRNA, encoding for no hairpin binding sites, without slncRNA induction (**E**) and with it (**F**). Both cells without NS induction and with it, present bright puncta localized to the poles. All scalebars are 5 µm. Color bars indicate mCherry fluorescence intensity.

To test the effect of hairpin-containing RNA, we designed a synthetic long non-coding RNA (slncRNA) sequence encoding for three hairpins to which QCP can bind, and cloned the corresponding DNA into the *E. coli* BL21 (DE3) strain under the control of a T7 promoter. Observation of cells in which protein expression is induced, but slncRNA expression is not (i.e., basal expression level dependent on the leakiness of the lacUV promoter controlling T7 RNA polymerase expression^45^) reveals homogenous fluorescence across the cells with no puncta whatsoever, indicating that low concentration of hairpin-containing slncRNA can effectively inhibit the phase-separating tendency of QCP (Figure 1C). In contrast, cells in which both slncRNA and protein expression are induced exhibit bright fluorescent puncta localized to the cell poles, indicating that an RNA-dependent heterotypic biomolecular condensate has emerged (Figure 1D). Similar expression of a negative control slncRNA (denoted as QCP-Non-Specific (NS)) which does not encode for any hairpins shows that polar puncta are retained, both with and without NS slncRNA induction (Figure 1E, F). To verify that our observations are indeed phase separation related and not an outcome of variations in protein content, we measured total cellular fluorescence using flow cytometry. The results, depicted in supplementary figure 1, show roughly equal fluorescence intensities for the various conditions appearing in figure 1, indicating that the localization and aggregation of the proteins is leading to the observed phenomenon.

### Synthetic RNP granules with Qβ coat protein exhibit polyphasic behavior

We hypothesized that RNA valency may be used to modulate the dissolution of the homotypic protein-only condensate and formation of heterotypic nucleoprotein structures in accordance with the polyphasic linkage model. To explore the effect of valency, we designed a set of 5 additional slncRNA sequences, encoding for 1,5,10,12, or 17 hairpins (denoted as QCP-*n*x with *n* being the number of hairpins encoded into the RNA sequence), and tested for the appearance of bright spots at the cell poles, with and without slncRNA induction. The results, presented as a fraction of spots per cell (Figure 2A), demonstrate different behaviors depending on slncRNA induction. Without induction (Figure 2A blue bars), bacteria expressing the QCP-NS slncRNA show the highest fraction of bright spots, as would be expected from a protein-based homotypic condensate. slncRNAs with low valency (QCP-1x/3x) appear to be sufficient to prevent the formation of the homotypic condensates but cannot phase separate on their own at low (uninduced) concentrations, resulting in virtually no visible spots. Bacteria expressing slncRNAs with higher valency (5 hairpins and above) show an abrupt jump to ratios in the range of 0.4 spots per cell. With induction (Figure 2A red bars), the spot-per-cell ratio increases as a function of valency, starting with an abrupt jump at a valency of three and peaking at a valency of ten, before subsequently decreasing for higher valencies.

**Figure 2.**
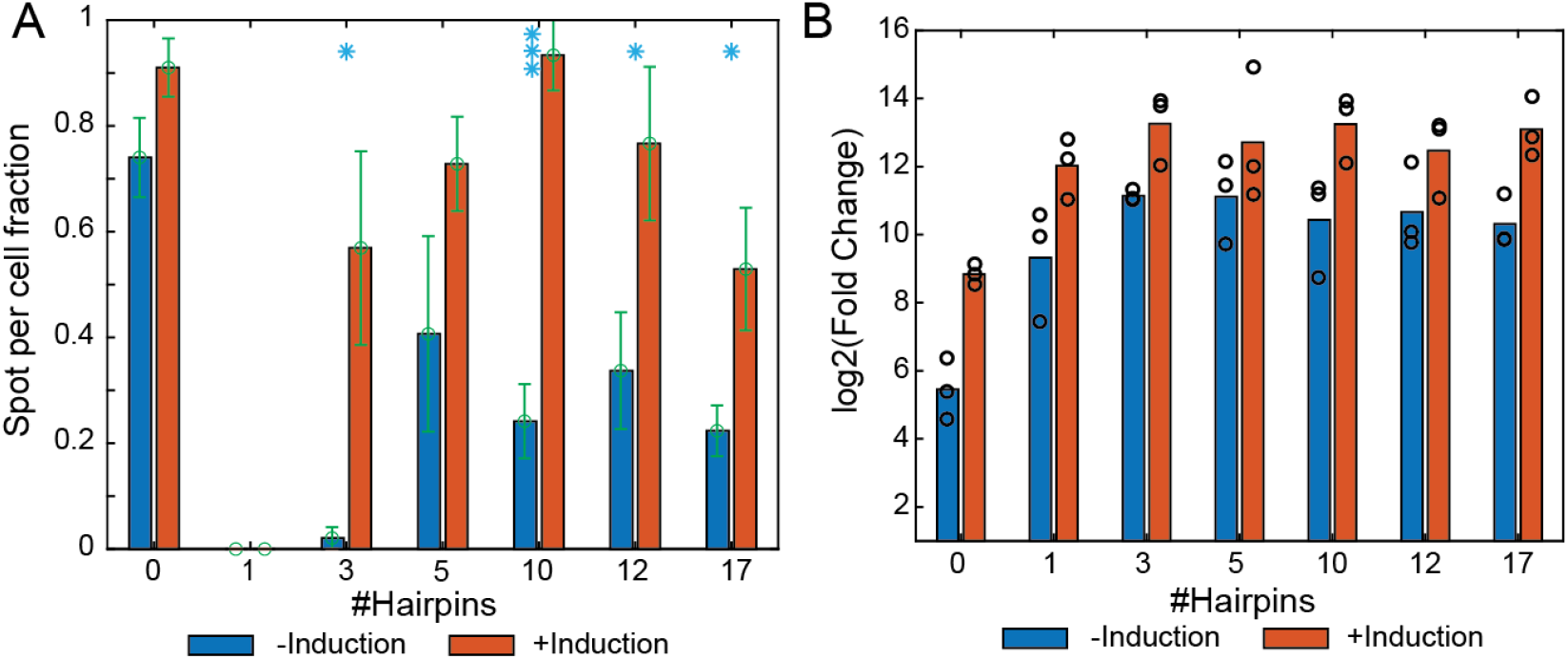
RNA valency and concentration can influence phase separation tendency. **A**. Spot-per-cell ratio for the different slncRNAs, with (red) and without (blue) induction of slncRNA expression. Ratio calculated as the number of fluorescent spots divided by number of cells, aggregated from multiple fields of view. Error bars indicate standard deviation. Stars indicate statistical significance (1 stars for the 0.05 threshold and 3 stars for the 0.001 threshold) according to two-sided t-test (p-values: QCP-3x: 0.03, QCP-10x: 9e-5, QCP-12x: 0.03, QCP-17x: 0.05). **B**. Log2(Fold Change) of the various slncRNAs at both induction levels, as measured via RT-PCR. slncRNA level per sample was quantified against the housekeeping gene idnT. Inter-sample slncRNA fold changes were calculated against the slncRNA levels of cells which do not contain a slncRNA expression vector (i.e., zero expression). The results show stable slncRNA expression levels for the various slncRNAs with a three orders of magnitude difference between the uninduced and induced cases.

To ensure that this observation is not a result of variations in slncRNA expression levels, we quantified these levels using RT-PCR. For this, the cells were grown in the presence of the appropriate inducers (50 µM C4HSL for protein induction and 0.33 mM IPTG for slncRNA induction), and cellular RNA was extracted following overnight growth. Expression levels for each slncRNA were measured against a known housekeeping gene *idnT*^46^. Fold changes in slncRNA expression were calculated relative to cells lacking a slncRNA expression vector. The results (Figure 2B) show that slncRNAs containing hairpins exhibit high abundance for n>=3, with roughly three PCR Cycles (∼x10) difference between uninduced and induced samples. By contrast, the QCP-NS slncRNA which does not encode for hairpins has low abundance in the cells with a nearly x32 reduced levels (∼5 cycles) independent of induction levels as compared with n>=3 valency slncRNA. A closer look reveals that the n=1 slncRNA system results in a significantly higher amount of RNA within cells as compared with the non-specific n=0 slncRNA (∼x16 or 4 cycles). This result together with the lack of phase-separation supports the notion that at n=1 a fully mixed state of nucleoprotein complexes forms, whereby the slncRNA is both protected from degradation and functions as a monovalent ligand which prevents the formation of homotypic protein-based biocondensates. Consequently, while these results verify that the differences in spot-per cell ratio do not originate from deviations in slncRNA abundance in the cells, they also provide further evidence for the existence of complex polyphasic behavior in our system.

### FRAP measurements provide support for four distinct states

To provide further support for the polyphasic behavior, we turned to a fluorescence recovery after photobleaching (FRAP) assay to probe the dynamics of the resulting condensates in the induced slncRNA (i.e. IPTG+) granule state^47^. To avoid the effects of condensate aging which could affect the results, we incubated the bacteria for 4 hours with the appropriate inductions (50 µM C4HSL for protein induction and 0.33 mM IPTG for slncRNA induction) prior to imaging. This duration was determined as optimal for both cell growth and condensate formation. To conduct the FRAP assay, we bleached the bright puncta and measured fluorescence recovery dynamics (Figure 3A). Fluorescence intensity quickly recovered (<15 sec) to some degree for slncRNA variants containing at least three hairpins, indicating a rapid exchange of proteins from the granules, while for the QCP-NS slncRNA lacking hairpins no discernible recovery was observed (Figure 3B). Further examination of the percentage of fluorescence recovery, also known as the mobile fraction, reveals two distinct features (Figure 3C). First, for valency of three and five hairpins, there is a sudden jump in the mobile fraction, from a non-recovered regime to a regime of substantial recovery. Second, at valency of ten hairpins or more, we observe a regime with smaller mobile fraction.

**Figure 3.**
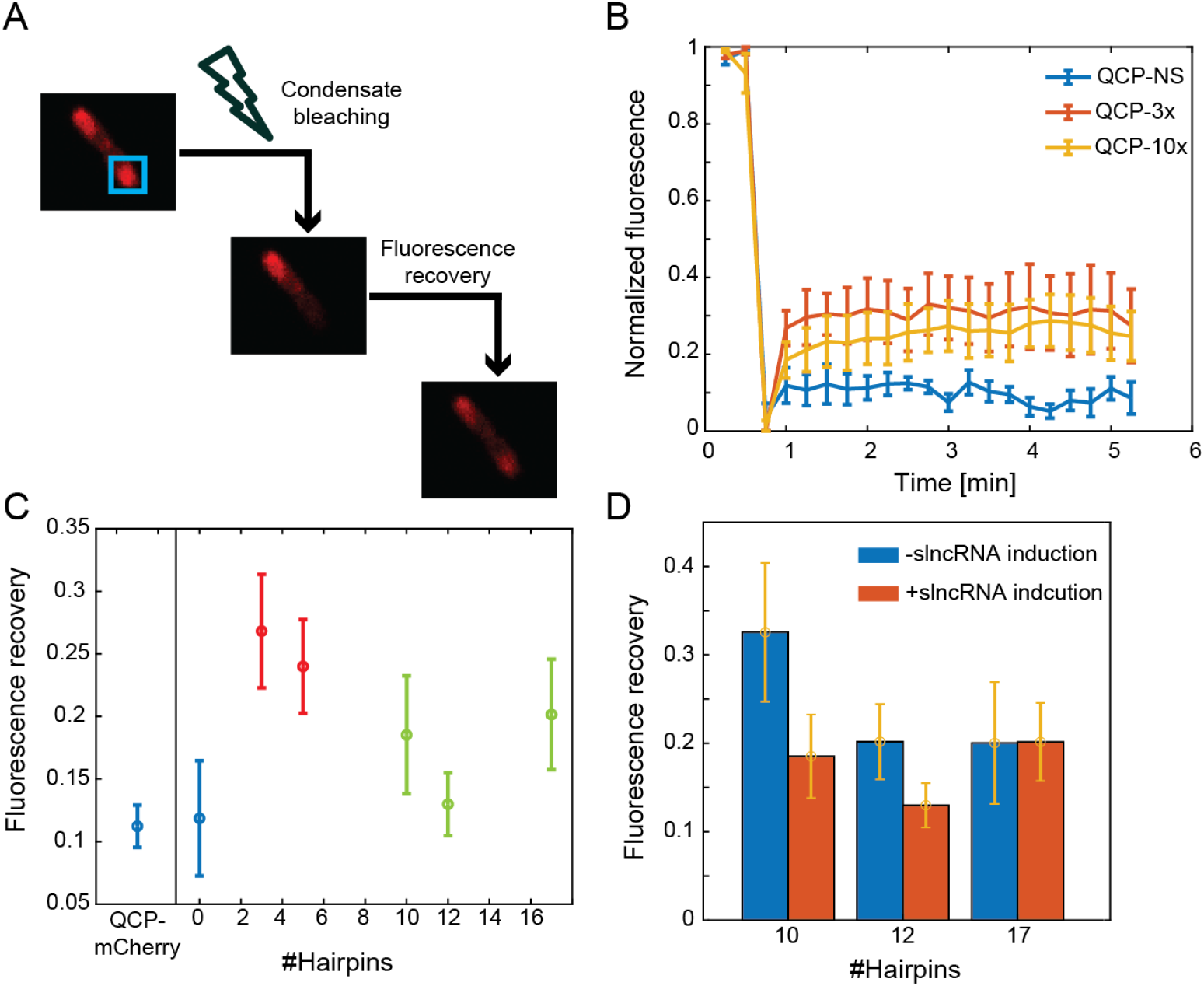
Fluorescence recovery after photobleaching reveals four different regimes. **A**. Fluorescence recovery after photobleaching (FRAP) pipeline. Bleaching was localized to a polar condensate, followed by temporal tracking to detect fluorescence recovery. **B**. Sample normalized fluorescence recovery curves measured from cells expressing the QCP-mCherry protein together with QCP-NS (blue), QCP-3x (red), or QCP-10x (yellow) slncRNAs. Curves present averages calculated from five different cells. Error bars indicate standard error of the mean. The curves show almost no recovery for QCP-NS, highest recovery for QCP-3x and intermediate for QCP-10x, representing low, intermediate, and high valency slncRNAs. **C**. Fluorescence recovery fractions (mobile fractions) for all tested slncRNAs. Condensates measured from cells which express the negative control slncRNA show similar average recovery to condensates which lack slncRNAs entirely, indicating a protein-dominant phase. Condensates measured from cells which express slncRNAs with three or five hairpins show highest recovery on average, implying a shift to a less-interconnected condensate with more molecular motion. Condensates measured from cells which express slncRNAs with more than ten hairpins demonstrate a drop in recovery fraction, indicating a more restrictive environment, probably due to RNA-RNA interactions becoming dominant. **D**. Fluorescence recovery fraction for QCP-10x/12x/17x with and without slncRNA induction. Condensate fluorescence recovers to a higher degree in the absence of induction, indicating more molecular motion. Bars represent mean values. Error bars indicate standard error of the mean. Data calculated from five separate cells.

We further measured the fluorescence recovery of granules formed with high valency slncRNAs (QCP-10x/12x/17x) without induction of slncRNA expression to observe the effects of slncRNA concentration. Here, the results (Figure 3D) show a mobile fraction that is either larger, or similar to, what was observed in the fully induced case, supporting a picture of different internal structure for the granules depending on RNA level. Together, the data shown in Figure 2 and 3 demonstrates at least four distinct biomolecular states: A homotypic protein-only condensate (NS), a mixed or non-interconnected nucleoprotein state (n=1,3), and apparently at least two heterotypic slncRNA-protein granular states. For lower valency (n<10) or slncRNA concentrations, the FRAP measurements suggest a granular state that is characterized by reduced structural interconnectivity as evidenced by the high mobile fraction. This reduced interconnectivity is reminiscent of a more “liquid-like” or “brittle” behavior, which can lead to an increased exchange of the biomolecules between the granulated state and the dilute surrounding environment. For higher valencies (n>10), the granules seem to be characterized by increased connectivity between the biomolecular components leading to a lower mobile fraction or more “solid-like” behavior. Consequently, these measurements indicate that two parameters, slncRNA concentration and valency, seem to allow control of internal interconnectivity and remodeling rate of the granules, which can in turn be utilized for different applications.

### QCP-mCherry titer supports a polyphasic dependence on slncRNA valency

A key feature of our previously reported synthetic RNP granules was their remarkable ability to increase the titer of the associated coat protein fusion inside the cell and form a type of protein-storage vessels, likely as a result of the granules protecting the protein component from cellular degradation processes. This hairpin-dependent increase in titer was visible in liquid culture and was estimated to be about 10-fold via flow cytometry measurements^36^. We hypothesized that a condensate with an RNA-binding protein containing an IDR would exhibit a more complex dependence of the protein titer on slncRNA valency. To test this, we measured total cellular fluorescence using flow cytometry of cells expressing QCP-mCherry together with the various slncRNAs, with and without induction of slncRNA expression, as well as cells expressing the mCherry protein alone (i.e., without the phage coat protein) together with the slncRNAs. To allow sufficient time for protein expression and subsequent granulation, we performed this measurement six hours after the introduction of the inducing molecules. The results (Figure 4A) demonstrate a substantial increase in fluorescence when comparing the QCP-mCherry fusion to the mCherry protein, regardless of slncRNA induction. The increase in fluorescence compared to mCherry alone is independent of induction level or valency of the slncRNA, and is consistent with previous reports by us ^48,49^ and others ^41^ that attribute this increase to the fusion to QCP.

**Figure 4.**
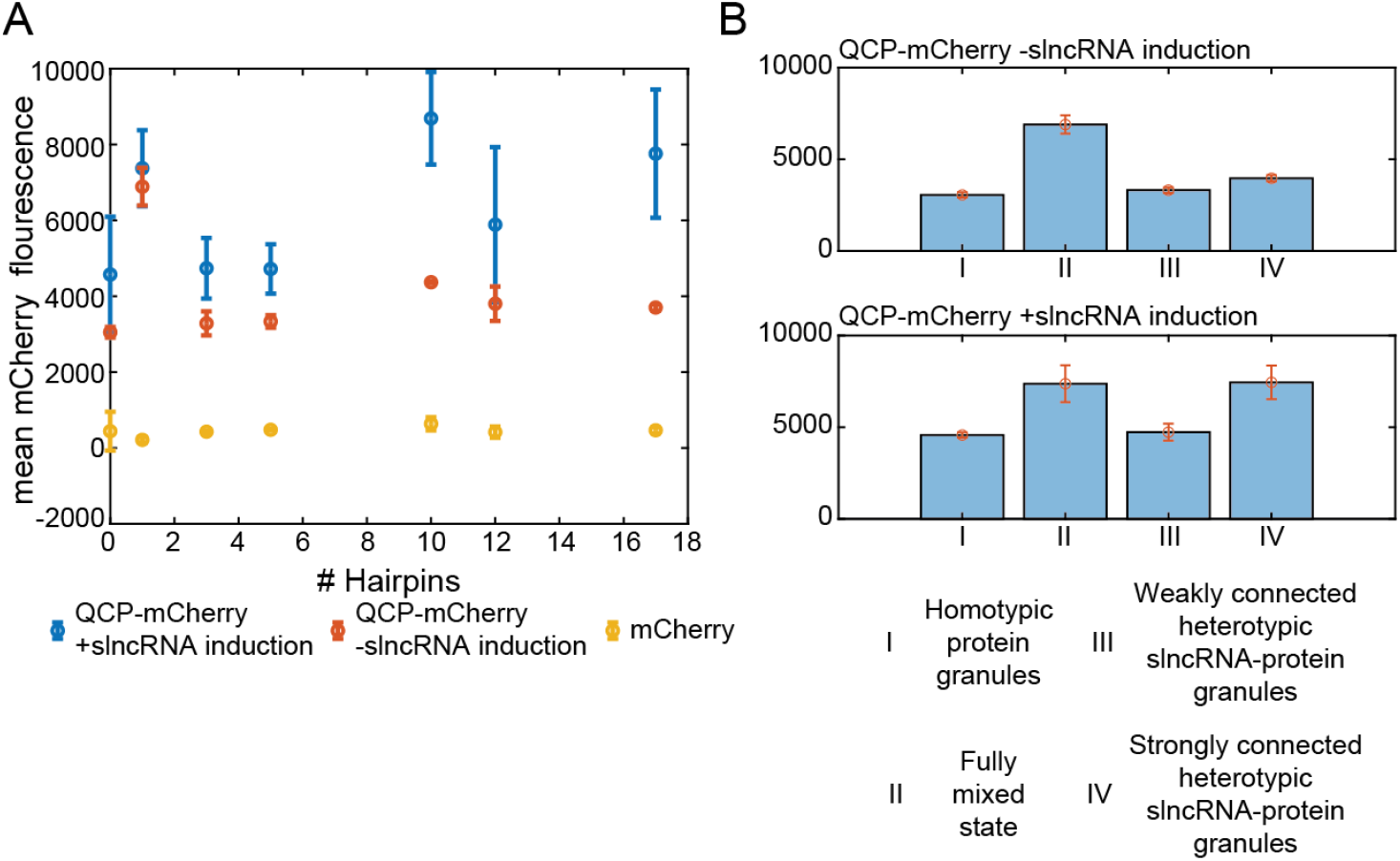
RNP granule solidity can affect protein titer. **A**. Mean mCherry fluorescence flow cytometry measurements from cells expressing slncRNAs together with QCP-mCherry with induction (blue), without it (red), as well as cells expressing slncRNAs together with the mCherry protein (yellow). The measurements show increased expression of QCP-mCherry and a non-monotonic relationship between fluorescence levels and hairpin number on the slncRNAs for the case of QCP-mCherry. Points indicate mean values; error bars indicate standard error of the mean. Values calculated from three biological repeats for all slncRNAs. **B**. Mean fluorescence measurements per state for QCP-mCherry without slncRNA induction (top), and QCP-mCherry with slncRNA induction (bottom), showing a dependence of fluorescence level on the internal state of the granules. Bars indicate average fluorescence; error bars indicate standard error of the mean.

A closer examination of the expression level data reveals a non-monotonic dependence of QCP-mCherry titer on valency for both the non-induced (red) and induced (blue) cases, which is consistent with polyphasic model supported by the cell fraction and FRAP measurements. Therefore, to further explore the fluorescence levels of QCP-mCherry under the assumption of the polyphasic linkage description, we aggregated the expression level of the four putative states and plotted the average fluorescence for each state per slncRNA induction level (Fig. 4b). For cells expressing the QCP-mCherry fusion without induction (Figure 4B - top), a strong increase in mean fluorescence of nearly x2 occurs between the homotypic protein-only granulated state (n=0) and the non-granulated fully mixed state (n=1). This is then followed by another x2 decrease to the weakly interconnected heterotypic slncRNA-protein granulated state (n=3 to 5). Then another potentially weak transition in fluorescence levels exists between the putatively weak interconnected state (mean fluorescence of 3.3e3 [A.U] for QCP-3x/5x), and the strong interconnected granulated state (mean fluorescence of 3.9e3 [A.U] for QCP-10x/12x/17x). While subtle, this shift is statistically significant, with a p-value of 0.01 using a two-sided t-test. In Fig. 4b-bottom we plot the mean fluorescence levels of the fully induced cells for all the different states. The data shows a similarly strong jump in fluorescence as for the non-induced state for the transition from homotypic to the mixed to weakly interconnected heterotypic states. However, the transition from the weakly to strongly interconnected heterotypic states is now more noticeable (mean fluorescence of 4.7e3 [A.U] for the weakly interconnected states slncRNAs, and 7.8e3 [A.U] for the strongly interconnected state, with a p-value of 0.025 using two-sided t-test). Consequently, the direct measurement of QCP-mCherry data provides further support for a multi-state model of our system as an RNA valency dependent transition from a homotypic protein only state to a mixed or melted state to a set of increasingly interconnected protein-RNA heterotypic granulated states.

### QCP granules increase protein titers for genes encoded on slncRNA cassettes

Given these results, we wondered how the protein titer of a gene encoded downstream to the slncRNA would be affected by the different states generated by this system as a function of slncRNA valency. To test this, we cloned a ribosome binding site and the gene for a blue fluorescent protein (TagBFP) downstream to our hairpin-encoding DNA sequence (Figure 5A). We first checked for the presence and localization of both the BFP and QCP-mCherry proteins using structured illumination super resolution microscopy and found that QCP-mCherry continued to aggregate at the cell poles while BFP fluorescence appeared evenly distributed throughout the entire cell (Figure 5B) as expected.

**Figure 5.**
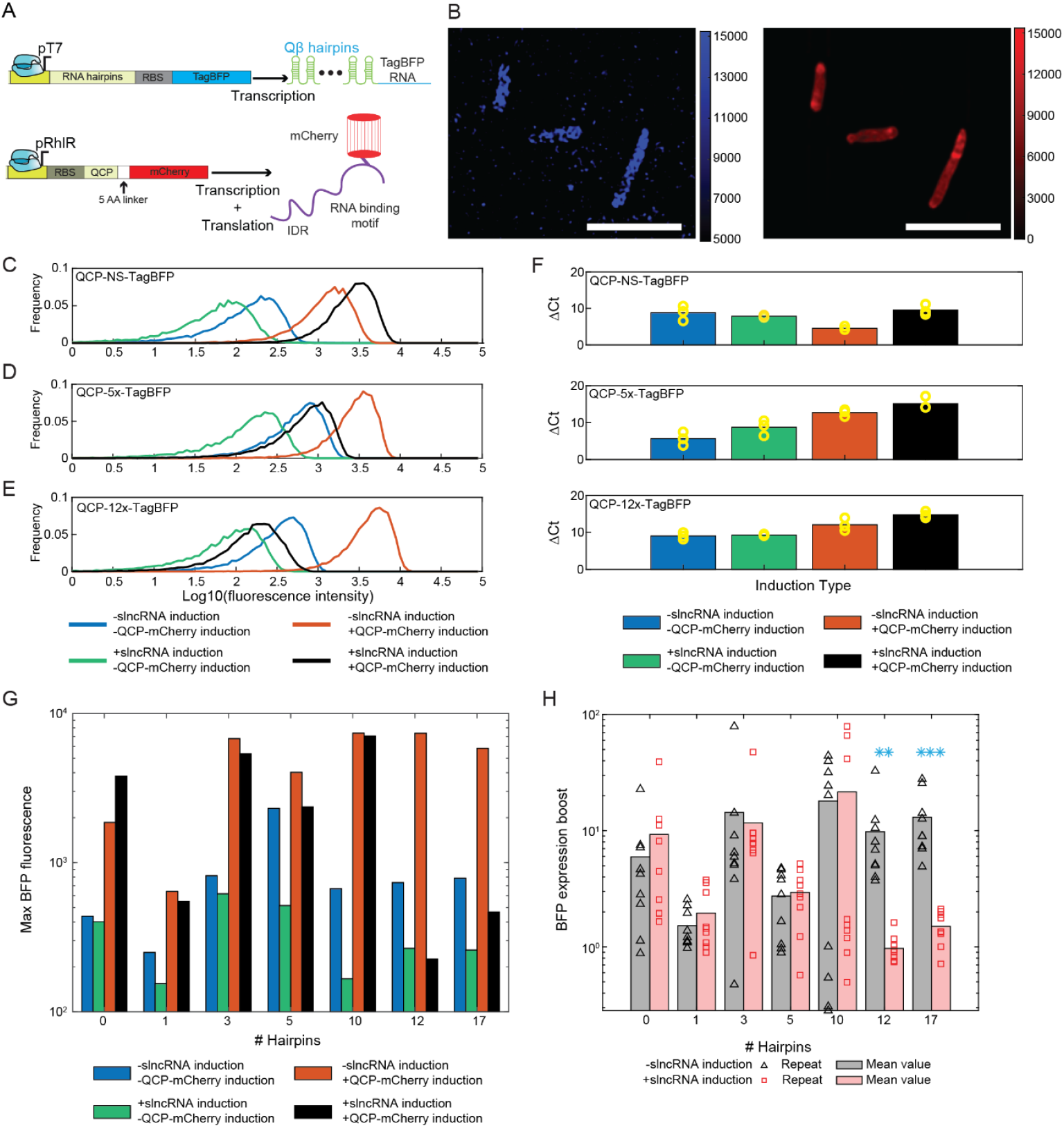
RNP granules can affect protein titer. **A**. Schematic of the genetic constructs: slncRNA-TagBFP under the control of a T7 promoter and QCP-mCherry under the control of a pRhlR promoter. **B**. Sample field of view showing cells with both QCP-mCherry (red) and TagBFP (blue) expression. Image captured using a structured illumination super resolution microscope. Scalebars are 5 µm. Color bars are in arbitrary units. **C-E**. Sample flow cytometry BFP fluorescence measurements of cells expressing QCP-mCherry together with QCP-NS-TagBFP (**C**), QCP-5x-TagBFP (**D**), or QCP-12x-TagBFP (**E**). **F**. slncRNA expression levels as measured in RT-PCR and presented as ΔCt values (calculated against the housekeeping gene idnT). The results show relatively constant expression of QCP-NS-TagBFP apart from one condition (the red bar corresponding to cells grown only with QCP-mCherry induction). QCP-5x-TagBFP and QCP-12x-TagBFP demonstrate an increase in slncRNA levels when QCP-mCherry is induced, indicating a protective influence of the condensates on the slncRNA. **G**. Maximal BFP fluorescence intensity values as a function of slncRNA valency and induction conditions. **H**. BFP expression boost defined as the mean BFP fluorescence intensity from cells with QCP-mCherry induction divided by the fluorescence intensity of cells without QCP-mCherry induction, Data is presented as a function of hairpins encoded on the slncRNAs. With slncRNA induction (red) or without it (black). Triangles and squares represent biological repetitions. Bars represent mean values calculated from all nine repeats. Stars represent statistical significance (2 stars for the 0.01 threshold and 3 stars for the 0.001 threshold) of a two-sided t-test (p values: QCP-12x-TagBFP: 0.01, QCP-17x-TagBFP: 7e-4.

We then measured BFP fluorescence via flow cytometry as a function of slncRNA induction, QCP-mCherry induction, and slncRNA valency. Figure 5C-E show representative fluorescence intensity distributions from cells expressing three different slncRNAs: QCP-NS-TagBFP, which does not encode for hairpins, QCP-5x-TagBFP, which encodes 5 RNA hairpins, and QCP-12x-TagBFP, which encodes 12 RNA hairpins, with all four combinations of induction. The results reveal a complicated dependency on slncRNA concentration and valency. Cells containing QCP-NS-TagBFP exhibit similar fluorescence levels without slncRNA induction, regardless of QCP-mCherry expression (Figure 5C blue and red traces). With slncRNA induction TagBFP levels increase in the presence of the QCP-mCherry protein (Figure 5C green and black traces), suggesting some influence of the protein condensate on the slncRNA. In contrast, QCP-5x-TagBFP and QCP-12x-TagBFP lead to more intricate behaviors. Cells grown without induction of either component exhibit intermediate levels of BFP fluorescence (Figure 5D, E blue traces), while those grown only with QCP-mCherry induction show the highest BFP fluorescence on average (Figure 5D, E red traces). Finally, cells grown with slncRNA induction demonstrate reduced fluorescence regardless of QCP-mCherry induction (Figure 5D, E green and black traces).

We wondered whether the differences in fluorescence intensity can be attributed to slncRNA expression levels. To investigate this, we quantified slncRNA expression using RT-PCR. The results, presented as ΔCt values (calculated against the housekeeping gene idnT ^46^), reveal that hairpin-containing slncRNAs show increasing expression levels across all the different induction conditions showing that increasing levels of QCP is associated with increasing levels of slncRNA-TagBFP. For these slncRNAs, the lowest expression is observed in uninduced cells (Figure 5F, middle and bottom plots, blue bars). Cells with slncRNA induction alone show a moderate increase in expression (green bars) from the non-induced cells, while cells with only QCP-mCherry induction (i.e. -slncRNA induction) display a noticeable jump in expression levels from the +slncRNA/-QCP-mCherry cells. The figures show jumps that range from 30 to 45% (i.e. 8.7 to 12.6 cell cycles for QCP-5x-TagBFP, and from 9.2 to 12.05 cell cycles for QCP-12x-TagBFP (red bars)). Finally, cells with full induction of both components show the highest expression levels (black bars) with an additional increase of ∼20% over the -slncRNA cells (black bars vs. red bars), and a total increase of 60-70% over the +slncRNA/-QCP-mCherry cells (black bars vs. green bars). These findings suggest that the condensates provide a protective environment for the slncRNAs, as indicated by their higher abundance in condensate-containing cells.

Interestingly, however, Fig. 5c-f also show that the increased abundance in slncRNA does not correlate with the fluorescence results. To fully map the relationship between valency and protein expression, we repeated the fluorescence measurements nine separate times for all slncRNAs and collected mean BFP fluorescence intensities. It is important to note that virtually all repeats demonstrated a single gaussian distribution, meaning that the entire population of cells (10k per measurement) shifted towards higher or lower fluorescence. In Fig. 5g we plot the maximal BFP expression levels for all the cassettes and induction levels as a function of their valency across the repeated growth experiments. The data shows that the highest BFP expression level is achieved for valency levels ranging from n=3 to 10, and for only QCP-mCherry induction, while fully induced cells with both high slncRNA and QCP-mCherry levels exhibit consistently lower expression level independent of valency.

To further evaluate this result, we calculated the boost in BFP fluorescence, which we define as the ratio between mean BFP fluorescence intensities of cells with QCP-mCherry induction, and cells without it. In Figure 5H we plot both the fluorescence boost of each individual repeat (black triangles for cells without slncRNA induction and red squares for cells with slncRNA induction) and the average value per slncRNA (black and red bars). The results display a wide range of values for each slncRNA, with some slncRNAs showing both a fluorescence decrease (<1x) and a massive increase (∼100x) in different biological repeats. In the range of n=1 to n=10 boost levels are consistent across the two different slncRNA induction levels. The lowest boost levels are observed for the n=1 valency consistent with it being characterized by a non-granulated mixed state. For n=3,5,10 higher values of boost are observed. Of particular note in that respect is QCP-10x-TagBFP, which shows a population of weak boost values and a separate population of extremely high values, both with and without slncRNA induction. For the n=12 and 17 cases, a distinctly different boost picture emerges. In both cases cells with high slncRNA concentration show practically no boost in BFP fluorescence, whereas the same cells with low slncRNA concentration present a moderately high increase. The difference in mean values within these slncRNAs is statistically significant with student t-test p-values of 0.01, and 7e-4 for QCP-12x-TagBFP, and QCP-17x-TagBFP, respectively. This result is consistent with nearly lack of BFP fluorescence observed for the fully induced cells in this valency range (Fig. 5g). Together these results provide further support for the polyphasic description of this system, and additionally indicate that the weakly interconnected heterotypic granule may be best suited for gene expression boost applications.

## Discussion

We studied the phase-separation behavior of synthetic RNA-protein granules, where the protein component (Qβ phage coat protein - QCP) contains both a specific RNA binding moiety and a known intrinsically disordered region (IDR). We characterized the QCP sRNP granules *in vivo* as a function of slncRNA valency (i.e., number of QCP binding sites) using cell fraction analysis, QCP-fluorescence intensity measurements, FRAP analysis, and via fluorescence measurements of a BFP gene encoded on the slncRNA.

Given the theoretical foundation of polyphasic linkage theory^34^ and our experimental data, we can construct a behavior profile for our system that is dependent on both RNA valency and concentration (Figure 6A). Non-valent RNA such as our QCP-NS results in granules dominated by homotypic protein-protein interactions, regardless of RNA concentration, implying that the Qβ coat protein acts as a scaffold molecule which undergoes phase separation. These granules are abundant in the cells, and exchange little to no material with the surrounding cellular environment. The presence of these protein granules boosts both QCP-mCherry and TagBFP titers (Figure 6A – regime I and Figure 6B), suggesting a potential influence of the granules on the RNA despite no apparent association between them. A possible explanation for this is non-specific interactions between the RNA component and the periphery of the already-formed protein condensates.

**Figure 6.**
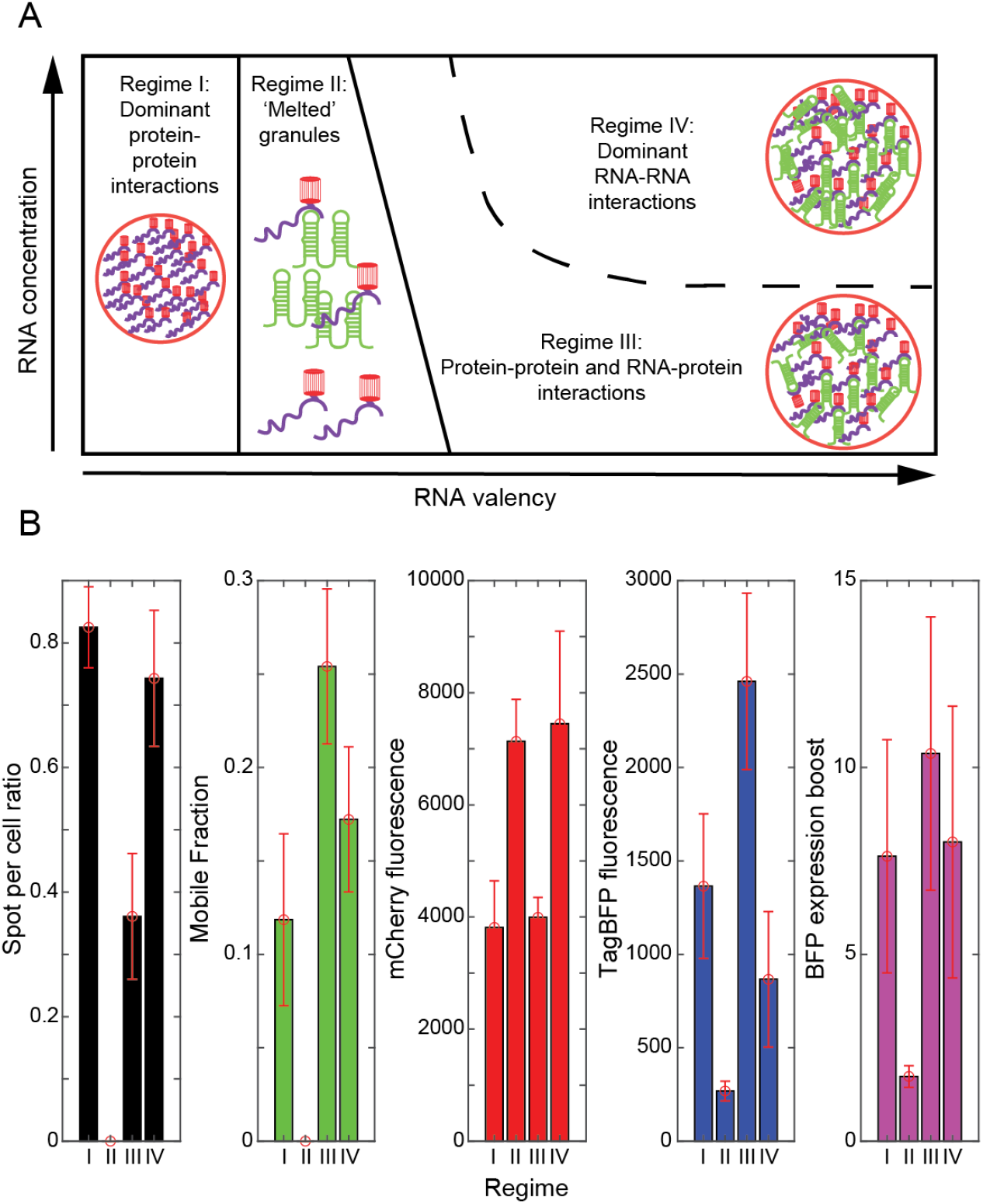
Internal granule regime can be determined by slncRNA valency and concentration. **A**. Granule biophysical characteristics can be described by four different regimes depending on RNA valency and concentration. Regime I: Non-valent RNA results in a protein-dominant regime with minimal exchange of molecules with the environment. Regime II: An increase in RNA valency leads to dissolution of the protein condensates, which we call the ‘melted’ state. Regime III: At intermediate valencies, granules are controlled by a combination of RNA-protein and protein-protein interactions, with maximal exchange of molecules with the environment. Regime IV: at high valency and high concentration, the granules become highly interconnected, with the RNAs likely functioning as scaffolds, and present lower molecular exchange. **B**. Aggregation of the statistical characteristics of each of the four described regimes as described in this paper: (from left to right) spot-per-cell ratio, mobile fraction, QCP-mCherry fluorescence, TagBFP fluorescence, and TagBFP expression boost.

At small valencies (n=1 to 3), we observe a regime with virtually no granules in the cells (i.e., the absence of spots for QCP-1x), which we refer to as the ‘melted’ or ‘non-interconnected mixed’ state. This phenomenon can also be explained by the polyphasic linkage model, where the proteins act as scaffold molecules which drive phase separation, and slncRNAs serve as ligands. QCP-1x, which can be considered a monovalent ligand, suppresses scaffold phase separation by either directly competing with the protein-protein interaction capacity of the coat proteins, or by enhancing their excluded volume ^34^. In this regime, QCP-mCherry fluorescence increases but BFP fluorescence is significantly reduced (Figure 6A – regime II and Figure 6B).

A further increase in the valency to intermediate levels (n=5 to 10) enables the formation of heterotypic protein-RNA interactions^50^. Here, the increase in spot-per-cell fraction indicates that protein-RNA interactions do not interfere with protein-protein interactions, thus contributing to the forces driving phase separation. According to the polyphasic linkage model, these intermediate valency slncRNAs function as multivalent ligands which bind to scaffold regions that do not actively promote phase separation (i.e., spacers). This augmented phase separation process leads to a more ‘relaxed’ internal environment which enables maximal exchange of molecules with the dilute phase (i.e., increased mobile fraction). This can then lead to the strong BFP expression and boost effect due to protection of the slncRNA from degradation leading to higher cellular titer, and sufficient “liquidity” allowing the slncRNA to access the cellular milieu and the available translation machinery with a small energetic cost (Figure 6A – regime III and Figure 6B). Finally, at high valency and high slncRNA concentration, the granules become dominated by RNA-RNA interactions leading to strongly interconnected structures. This results in a substantial slowing down of the macromolecular exchange between the granule and the cellular milieu as evidenced by the smaller mobile fraction, which leads to a significant drop in TagBFP expression (Figure 6A – regime IV and Figure 6B). It is important to note that while polyphasic linkage theory provides an adequate qualitative model for describing the transition between the weaker and stronger interconnected states for higher valencies, the actual biophysical mechanism which underlies this phenomenon (e.g. transition from an IDR-dominated association to a Percolation-dominated state) remains to be resolved.

From a technological perspective, these results indicate that the QCP granule platform may be used as a tool for increasing the titer of protein production within bacteria. Specifically, our findings suggest two different approaches. The first relies on fusing the target protein to QCP. Here, we found a maximal boost of ∼30x on average for mCherry fluorescence using the n=1 valency slncRNA as compared with the case for mCherry alone. The second approach is to encode a gene directly downstream from the protein binding moiety. This approach yielded a boost of up to x80 in BFP expression for the n=10 valency. Nevertheless, several challenges remain when considering biotechnological applications using either approach. Notably, for the first approach the protein mass is potentially trapped inside a granule in which the correct folding of proteins is possible, but not guaranteed. Furthermore, efficient extraction of the protein from the granules might require heating or use of other denaturing agents, potentially damaging or destroying the target protein in the process. Finally, the extracted protein is a fusion between the desired protein and QCP, requiring protease cleavage and a subsequent purification step to obtain a final product. For the second approach, a lack of reproducibility was observed, where some biological repeats yielded a large boost factor, and others of the same system exhibited a much different result, with particularly sharp dichotomy observed for the n=10 valency slncRNA. This could be a result of internal factors such as sub-optimal slncRNA design, which leads to the protein-protein interactions outcompeting the slncRNA gel interconnected phases (in other words, the protein-protein interactions of the IDR overcome the protein-slncRNA interactions), or granule expressing cells and their internal organization may be more sensitive to external factors such as small differences in oxygen supply due to different positions within the incubator. Additionally, slncRNA valency appears to be a critical factor for this approach, as high valency leads to reduced expression, likely because of lower accessibility to the translation machinery. Consequently, additional study is needed to convert these findings to a robust biotechnological application.

Finally, recent works on synthetic phase separation systems have focused on intrinsically disordered elements as the drivers of phase separation^51,52^. To the best of our knowledge, this work constitutes a first-of-its-kind completely synthetic model of naturally occurring RNP granules in which both RNA and protein components not only contribute to the formation of the granules but are also capable of changing internal granule features and dynamics. We have demonstrated that RNA valency with respect to the protein can overcome protein-protein interactions mediated by intrinsically disordered regions and lead to a nearly complete dissolution of the protein granules in low valency RNA, or to the formation of an RNA-dominant granule. The concept of IDR modulation via RNA, as demonstrated here, could be a promising therapeutic strategy for pathological conditions related to changes in IDR behavior ^53^. Finally, the synthetic system presented here could be a useful tool to further explore polyphasic as well as multiphasic condensate formation provided that naturally occurring IDRs are included enabling a more controlled exploration of complex phase behaviors.

## Data availability

The data underlying this article is available in the article.

## Acknowledgements

We thank Dr. Nitsan Dahan and Dr. Yael Lupu-Haber from the Technion Life Science and Engineering infrastructure center for their support in microscopy experiments.

## Author contributions

N.G. and R.A. conceived and designed the research; N.G. and S.G. performed the molecular biology cloning steps; N.G. performed the experiments and data analysis; N.G. and R.A. wrote the paper.

