## Supplementary information for "Formation of polyphasic RNP granules by intrinsically disordered Qβ coat proteins and hairpin-containing RNA"

**Authors**: Naor Granik[1], Sarah Goldberg[2] and Roee Amit[2,3*]

**Affiliations**:

1 Department of Applied Mathematics, Technion - Israel Institute of Technology, Haifa 32000, Israel.

2 Department of Biotechnology and Food Engineering, Technion - Israel Institute of Technology, Haifa 32000, Israel.

3 The Russell Berrie Nanotechnology Institute, Technion - Israel Institute of Technology, Haifa 32000, Israel.

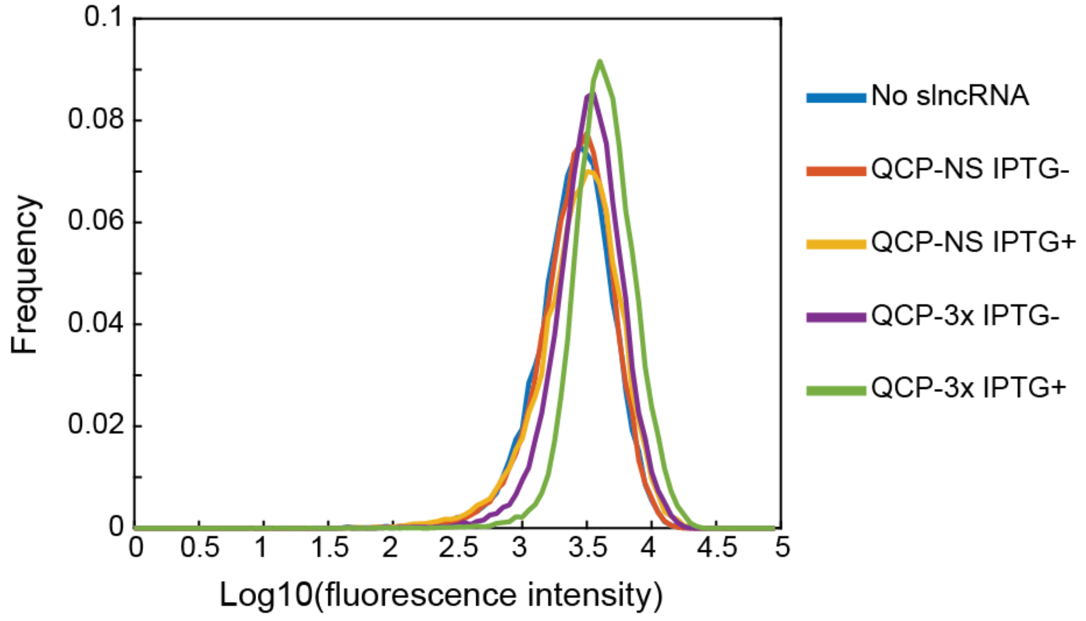


Supplementary Figure 1.

Flow cytometry measurements of E. Coli BL21 expressing QCP-mCherry alone (blue trace) or together with QCP-NS (uninduced – red trace; induced – yellow trace) or QCP-3x (uninduced – purple trace; induced – green trace). The measurements show roughly equal fluorescence levels between the different conditions, indicating similar levels of protein content
